# Validation of a colorimetric LAMP to detect *Loxosceles* experimental envenomation

**DOI:** 10.1101/2022.02.09.479769

**Authors:** Luana Paula Fernandes, Marcele Neves Rocha, Clara Guerra Duarte, João Carlos Minozzo, Rubens L. do Monte-Neto, Liza F. Felicori

## Abstract

Diagnostic tests for brown spider accidents are unavailable and impact treatment decisions, increasing costs and patient risks. In this work, we used for the first time a fast, simple, and visual method based on the loop-mediated isothermal amplification assay (LAMP) to detect *Loxosceles* envenomation. Using the DNA from *L. similis* legs, we observed a high sensitivity using this test since as low as 0.32 pg of DNA could be detected. This pH-dependent colorimetric assay was 64 times more sensitive than PCR to detect spider DNA. The test was specific for *Loxosceles* once no cross-reaction was observed when testing DNA from different agents that cause similar dermonecrotic injuries. The test allowed the detection of *Loxosceles intermedia* DNA from hair, serum, and swab samples obtained from experimentally-envenomed rabbit within 72 h. The method sensitivity varied according to the sample and the collection time, reaching 100% sensitivity in serum and hair, respectively, 1 h and 24 h after the experimental envenomation. Due to its ease of execution, speed, sensitivity, and specificity, LAMP presents an excellent potential for identifying *Loxosceles* spp. envenomation. This can reduce the burden on the Health System and the morbidity for the patient by implementing the appropriate therapy immediately.In addition, this work opens up the perspective to other venomous animal accident identification using LAMP.

**Highlights:**

* Using 28S primers it was possible to identify *L. similis*’ DNA with high sensitivity;
* LAMP was 62-fold more sensitive than PCR and detected as low as 0.32 pg of DNA;
* LAMP detected *L. intermedia* DNA from hair, serum, and exudate from experimentally-envenomed rabbits;
* LAMP presents an excellent potential for identifying *Loxosceles* spp. envenomation.

## 1. Introduction

Accidents with spiders of the genus *Loxosceles* (brown spider) represent a serious global health problem, mainly due to the morbidity associated with the bite of these animals. The clinical manifestations of these accidents, known as loxoscelism, are considered the most serious among the spider genera. The characteristic of the cutaneous form of loxoscelism is a dermonecrotic lesion that is hard to heal, accompanied by nonspecific systemic symptoms such as nausea and fever [1–3]. In the cutaneous-visceral loxoscelism, dermonecrotic lesion is usually accompanied by vascular manifestations, such as hemolysis (intravascular or extravascular), which can progress to acute renal failure and, in some cases, to disseminated vascular coagulation considered as the leading cause of death from loxoscelism [4,5].

The early identification of this envenomation makes it possible to use specific treatments such as serum therapy to prevent the progression of systemic symptoms resulting from loxoscelism. However, this identification occurs in less than 20% of the reported cases, mainly because the bite is not painful and goes unnoticed by the victim. In addition, in the beginning, the lesion can be, for example, misdiagnosed as bacterial infection (*Staphylococcus*, *Mycobacterium*, *Syphilis*, *Pseudomonas*, *Rickettsias*), fungal infection (*Sporothrix schenckii*), viral infection by herpes, leishmaniasis, diabetic ulcers, erythema nodosum and Lyme disease[3,6]. Facing this scenario, it is necessary to develop a quick and simple method that allows the precise differential diagnosis of loxoscelism. Studies have already explored ELISA-based techniques (sandwich and competition) to detect protein components of *Loxosceles* venom in animal samples. However, these techniques are time-consuming and present low sensitivity (reviewed by [7]. To increase sensitivity and specificity, DNA-based identification methods, such as PCR (polymerase chain reaction), could also be an alternative, as it has already been pursued to snake envenomation [8,9] However, PCR requires well-trained staff and bench thermocyclers, limiting its use in the field and resource-limited areas. Portable and miniaturized devices, though, can be an alternative for that. The loop-mediated isothermal amplification (LAMP) method is cost-effective (1.50 USD/test), simple (the isothermal reaction requires a simple heating device), fast (results within 60 min) [10], and visually detected [11]. Because of this, LAMP has been used to detect parasites, [12–15], bacterias, [16,17], sexually transmitted diseases [18–20], and viruses including SARS-CoV-2 [21–23].

Therefore, in this study, LAMP was applied to detect *Loxosceles intermedia* DNA in serum, exudate, and hair samples collected from experimentally-envenomed rabbits. This is a pioneer study devoted, for the first time, to detect DNA from venomous animals envenomation, different from previous studies where venom protein components or antibodies against venom were evaluated.

## 2. Materials and methods

### 2.1. Animals and venoms

*Loxosceles* spider’s crude venom - obtained from *L. intermedia*, *L. gaucho*, and *L. laeta* - were collected and provided by the Production and Research Center of Immunobiological Products (CPPI), Paraná, Brazil. Ten-week-old New Zealand female rabbits were experimentally envenomed with 5 μg of *Loxosceles intermedia* spider venom, followed by extraction of samples, such as blood, hair, and exudate. Animals were maintained in the animal facility at the Instituto de Ciências Biológicas - Universidade Federal de Minas Gerais (UFMG) and received water and food *ad libitum*. Treatment and handling of all animals used were in accordance with the Ethics Committee on the Use of Animals (CEUA)/UFMG, license number 291/2019. The spider *Loxosceles similis* was collected in the city of Nova Lima, Minas Gerais, Brazil, under authorizations of the Brazilian Authorization and Biodiversity Information System (SISBIO) (Process number 72083-1), and the National System for Management of Genetic Heritage and Associated Traditional Knowledge (SISGEN) (Process number 72083-1) by the Fundação Ezequiel Dias (FUNED) in Belo Horizonte. The spider was kept at 24 °C and fed weekly with crickets until its use.

### 2.2. DNA extraction Obtention of DNA from *Loxosceles similis* legs

DNA was extracted from the legs of the spider *Loxosceles similis* and used as positive control. The DNA extraction was performed as recommended by the manufacturer using the QIAamp DNA Extraction Reagent Kit (Qiagen).

### 2.3. LAMP primer design

LAMP primers were designed to target the 28S ribosomal RNA gene from spiders belonging to the genus *Loxosceles* (Access in GenBank No. EU817786.1). The 28S consensus sequences among *L. laeta*, *L. gaucho*, *L. intermedia*, *and L. hirsuta* were aligned using MUSCLE, [24]. The regions with low or no identity with sequences from organisms that misdiagnose with *Loxosceles* (*Mycobacterium tuberculosis*, *M ulcerans*, *Staphylococcus aureus*, *Syphilis treponema*, *Rickettsia rickettsii*, *Pseudomonas aeruginosa*, *Chromobacterium violaceum*, *Sporothrix schenckii*, *Aspergillus sp*, *Cryptococcus sp*, *Leishmania sp*, *Herpes simplex*) were considered [3,6,25,26]. Six LAMP-specific primers (two internal – FIP and BIP, two external – F3 and B3, and two loop primers – LF and LB) were then generated using Primer explorer V5 (http://primerexplorer.jp/lampv5e/index.html) and analyzed using Multiple Primer Analyzer from Thermo Scientific. The oligos were purchased from GenOne and validated using *L. similis* DNA as template (Table 1).

**Table 1:**
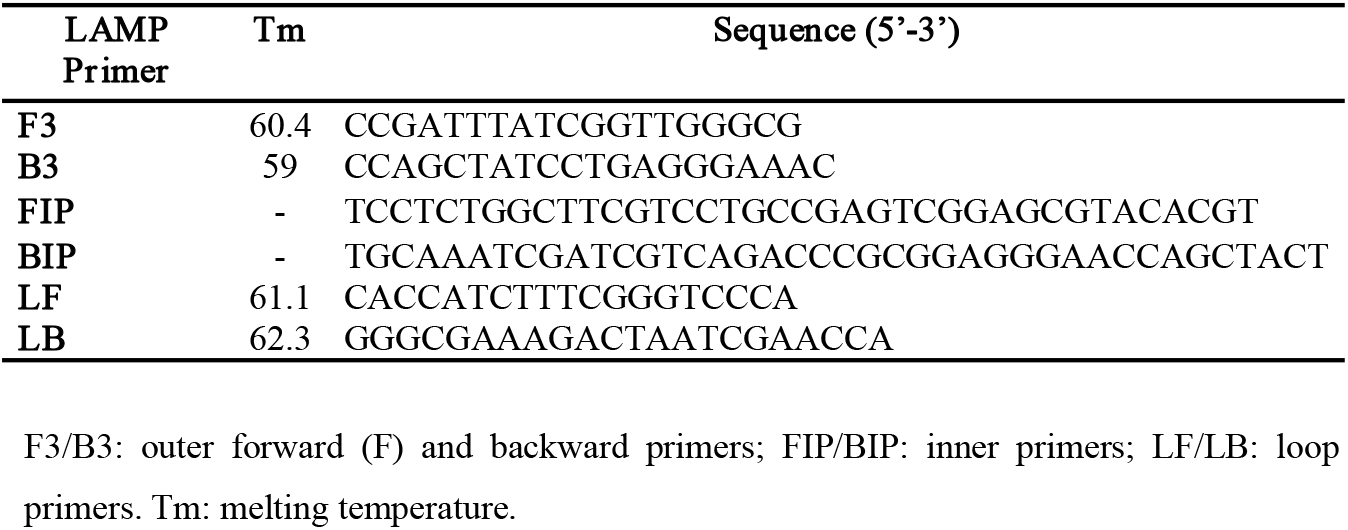
Set of LAMP oligonucleotides designed and used in this study

### 2.4. LAMP assay

LAMP assays were optimized with *Loxosceles similis* DNA using Master Mix reagent (WarmStart^®^ #M1800 – New England BioLabs). For this, different settings were tested concerning: the primer concentration (F3/B3: 0.05 to 0.2 μM; FIP/BIP: 0.4 to 1.6 μM and LF/LB: 0.1 to 0.4 μM); temperature (60, 65, 68 or 71 °C) and the amplification time (15, 30 or 60 min). The amplification products were analyzed by resolving them in 1.5 % agarose gels and visually monitored since the reaction buffer contains the pH indicator phenol red that turns from pink to yellow. Gel images were taken using the L-Pix Chemi Molecular Imaging.

### 2.5. PCR assay

Conventional PCR was also performed to compare the sensitivity and specificity of *Loxosceles* DNA detection in the collected samples. For this, 1.5 μL of PCR buffer, 1.5 mM of MgCl_2_, 0.2 mM of dNTP mix, 1.2 μM of forward external primer (F3), 1.2 μM of backward external primer (B3), 1 unit of Taq platinum DNA polymerase enzyme was added to a final volume reaction of 25 μL in water. PCR programming consisted of initial denaturation at 94 °C for 2 minutes, followed by 35 cycles at 94 °C for 30 s, (55 or 60 or 65 °C) for 30 s and 72 °C for 1 minute, followed by 5 min final extension. The amplification took place in a thermocycler (SimpliAmp Thermal Cycler – Thermo Fisher). The amplification product was evaluated on agarose gels (1 % w/v) with SYBR safe (0.009 % v/v) (Invitrogen) in 1X Tris/Borate/EDTA buffer (TBE).

### 2.6. Limit of detection (LoD), sensitivity and specificity

The sensitivity and specificity for LAMP with DNA from *Loxosceles similis* was evaluated using the best LAMP conditions: 0.2 μM F3/B3, 0.4 μM LF/LB, 1.6 μM FIP/BIP primers; at 71 °C incubated during 60 min. For sensitivity, *Loxosceles similis* DNA was titrated (10 ng, 1 ng, 0.1 ng, 0.01 ng, 0.005 ng, 1.5 pg, 1.25 pg, 0.62 pg, 0.32 pg, 0.15 pg) and used as the template. To PCR assays, the input of *Loxosceles similis* DNA were: 10 ng, 5 ng, 1 ng, 0.5 ng, 0.1 ng, 0.02 ng, 0.01 ng and 0.002 ng. For specificity evaluation, five microorganisms were used: *Leishmania braziliensis*, *Herpes simplex virus* (*HSV-1*) *Rickettsia rickettsii*, *Rickettsia parkeri* and *Corynebacterium pseudotuberculosis*. In addition, the specificity of the negative control samples (collected prior to the experimental rabbit envenomation) was also evaluated. These different microorganisms’ DNA was obtained in partnership with Fundação Oswaldo Cruz - Fiocruz, Brazil. DNA from negative controls was obtained from sample extraction performed with the phenol:chloroform;isoamyl alcohol method, because it is effective and allows large-scale extraction. The concentration of DNA total obtained varied between 11-20 ng/μL (*swab*), 17-24 ng/μL (*hair*), and 130-260 ng/μL (*blood*). The influence of serum and saline on LAMP reactions was evaluated since these components are present in blood and hair samples, respectively.

### 2.7. Experimentally-envenomed rabbit samples processing

Serum, exudate, and hair samples were obtained by experimental envenomation of 6 rabbits weighing between 1.5 and 3 kg. The animals were inoculated with 5 μg of *Loxosceles intermedia* spider venom. Samples were collected before (negative control) and after envenomation at 1, 8, 24, 48, 72, and 240 h. Exudate samples were collected using cotton swabs. The swab was immersed in 0.9 % saline solution and kept over the venom inoculation area for 30s. The samples had the DNA extracted using the phenol:chloroform:isoamyl alcohol method. For the extraction, 500 μL serum, 500 μL saline (swabs), and 30 hair bulbs were used for each collected sample. Thus, to each sample was added: 300 μL TNE buffer (50 mM Tris-HCl, 100 mM NaCl, 6.3 mM EDTA, pH 7.5); 10 μL proteinase K solution (10 mg/mL); 7 μL CaCl_2_ (0.5 mM); 10 μL SDS (25 %) and 100 μL 2-mercaptoethanol, homogenized and incubated at 55 °C for 3 h. Then, 300 μL phenol:chloroform:isoamyl alcohol (25:24:1) was added and centrifuged 10,000 *g* for 15 min. The supernatant was transferred to a new tube followed by the addition of 300 μL absolute ethanol (−6 °C) and 50 μL 3 M sodium acetate pH 5.2. It was centrifuged again at 10,000 *g*. The pellet was washed with absolute ethanol twice. After drying at room temperature, the DNA was diluted in 50 μL milli-Q water. After extraction, DNA was quantified using NanoDropTM One/One^c^ (Thermo Fisher Scientific).

### 2.8. Statistical analyses

Comparative analyses were performed in relation to negative controls at each collection time point (1, 8, 24, 48, 72, and 240 h). Six rabbits were included per group (time point). The variables were qualitative (positive and negative) and paired samples (negative controls and samples taken from the same animal) were used. Thus, the statistical test used for the analyzes was the non-parametric Chi-square. The test’s sensitivity (positive results among those envenomed) and specificity (negative results among those not envenomed) were also evaluated for each sample at different time-points.

## 3. Results

### 3.1. *Loxosceles similis* DNA can be detected by LAMP, which is 62-fold more sensitive than PCR

The best LAMP condition selected and used in all of the following reactions were: 0.2 μM F3/B3, 0.4 μM LF/LB, 1.6 μM FIP/BIP primers; at 71 °C incubated for 60 min (Supplementary Figure S1). Using this condition, the observed LoD, which is the lowest detectable *Loxosceles* DNA concentration, was 0.32 pg. This was confirmed by visual colorimetric LAMP and by resolving the amplified DNA in agarose gel (Figure 1A). The LAMP sensitivity was 62-fold higher than that of PCR, detecting 0.02 ng of *L. similis* DNA (Figure 1B).

**Figure 1.**
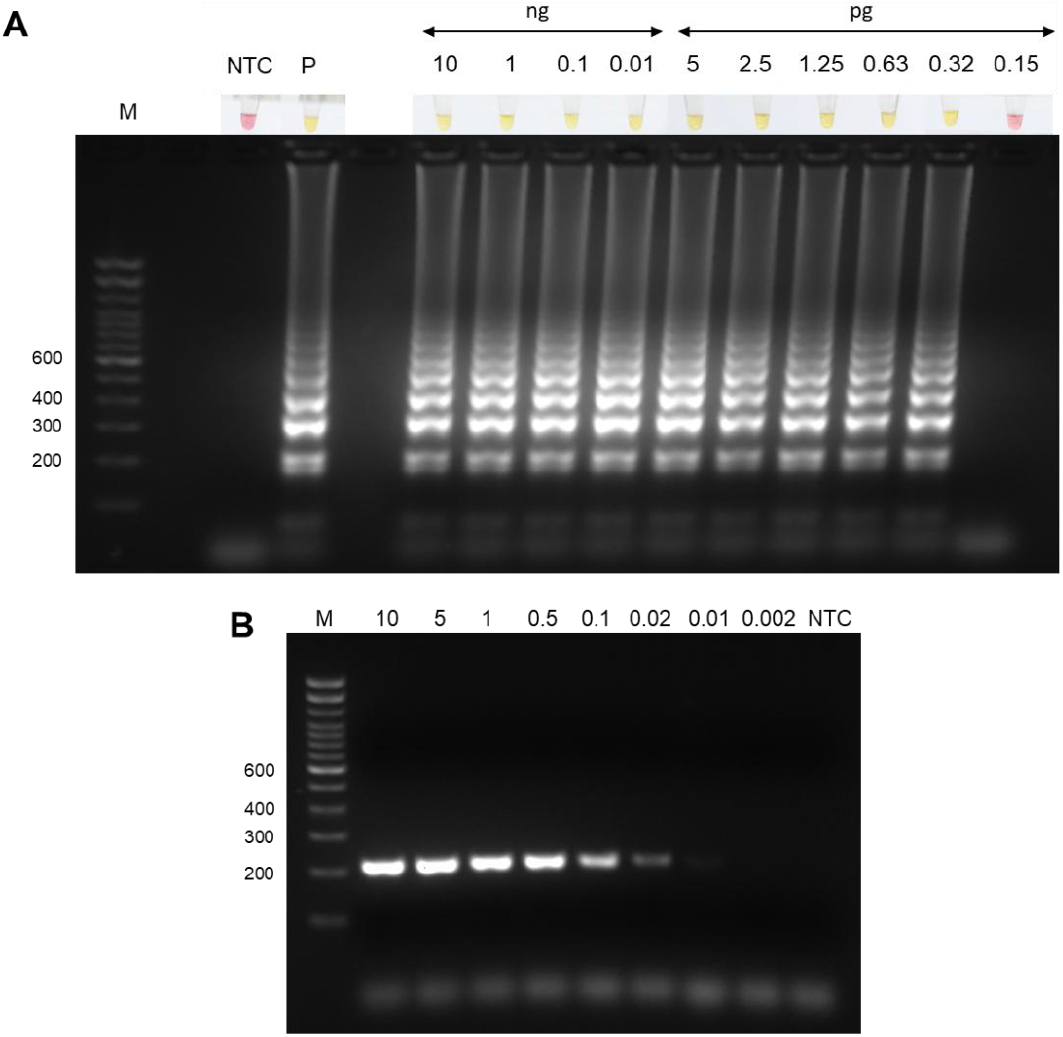
Limit of detection of *Loxosceles similis* DNA with LAMP and PCR. A) LAMP reaction was performed at 71 °C for 60 min using WarmStart^®^ colorimetric master LAMP mix (NEB #M1800) in 20 μL final volume. Amplicons were resolved in 1.5% agarose gel and stained with SYBR safe (0,009% v/v) (Invitrogen) to confirm DNA amplification. The LoD was established by titrating the *L. similis* DNA as input, ranging from 10 ng to 0.15 pg. B) PCR amplicons obtained with different *L. similis* DNA inputs varying from 10 to 0.002 ng. The assay was performed with external primers (F3 and B3) and TaqPlatinum™ enzyme. M: molecular weight standard of 100 bp. NTC: no template control, P: Positive control (10 ng *L. similis* DNA).

### 3.2. LAMP is specific for *Loxosceles similis* DNA

We also observed that the LAMP assay was specific for the *L. similis* DNA when tested with other organisms DNA that would cause dermonecrotic as a clinical manifestation similar to loxoscelism, such as *Rickettsia rickettsii*, *Rickettsia parkeri*, *Leishmania braziliensis*, *Corynebacterium pseudotuberculosis*, and *Herpes simplex virus-1* (Figure 2A). *PCR was also specific for L. similis DNA* when the other samples were tested (Figure 2B). We also demonstrated that it is possible to identify the spider’s DNA in the *Loxosceles* crude venom (Supplementary Figure S2).

**Figure 2.**
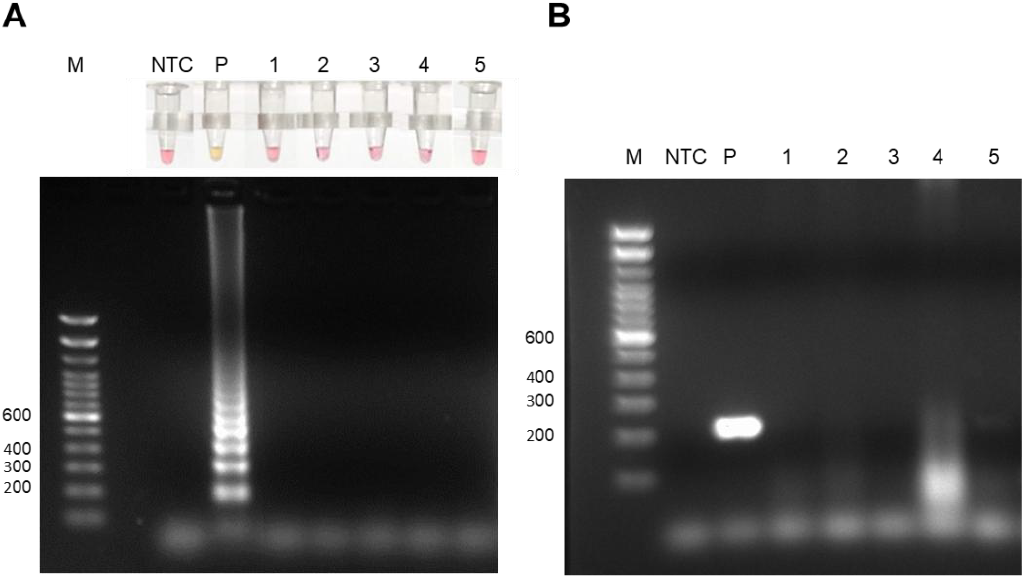
Specificity for the detection of *Loxosceles similis* DNA using LAMP or PCR. 20 ng of DNA from *Rickettsia rickettsia* (1), *Rickettsia parkeri* (2), *Leishmania braziliensis* (3), *Corynebacterium pseudotuberculosis* (4), and *Herpes simplex virus-1* (5) were tested using LAMP (A) or PCR (B). M: Molecular weight marker. NTC: no template control. P: Positive control (10 ng *L. similis* DNA).

### 3.3. Detection of *Loxosceles* DNA in rabbit samples by LAMP assays

We were able to detect *Loxosceles* DNA in all samples (serum, exudate, and hair) collected from 6 different rabbits before and after experimental envenomation in six different time points. Their DNA was extracted and then evaluated by LAMP. For hair and serum samples (Figure 3A and Figure 3C), detection could be observed from 1 to 72 h, while detection in exudate samples was observed up to 24 h after envenomation (Figure 3B). We can also observe that 100 % sensitivity was observed in serum samples 1 h and in hair samples 24 h respectively (Figures 3A and 3C, Table 2).

**Figure 3.**
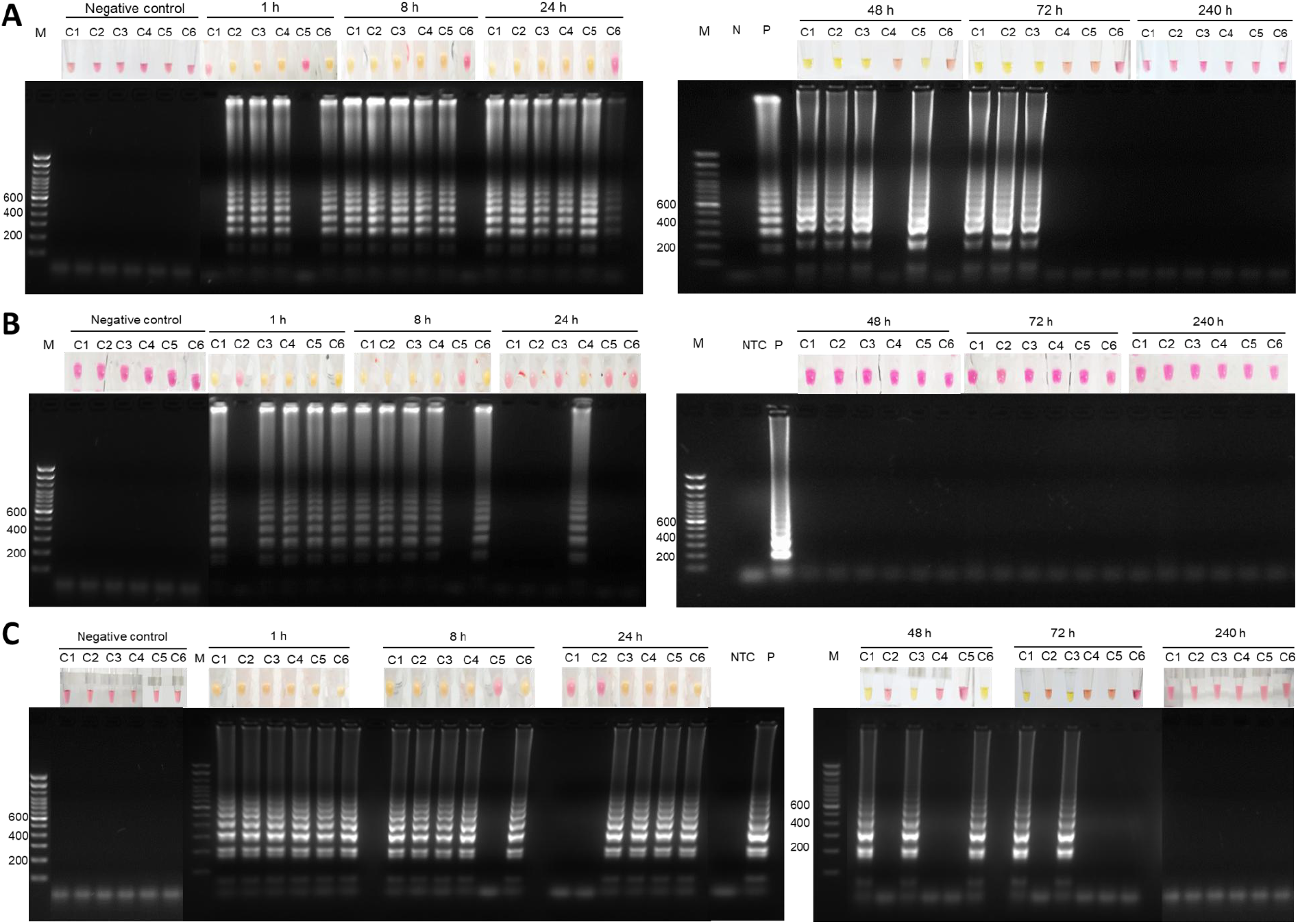
Detection of *L. intermedia* DNA in hair (A), exudate (B) and serum (C) samples from experimentally envenomed rabbits by LAMP. Colorimetric output for each sample within six times after envenomation. 1.5 % agarose gel for time samples. M: molecular weight marker. Negative control: samples collected prior to poisoning. NTC: no template control. P: positive control (10 ng *L. similis* DNA). C1 to C6 refers to rabbits 1 to 6.

**Table 2:** LAMP sensitivity for detecting different rabbit samples in different time-points.

| Sample type | Time points (h) |  |  |  |  |  |
| --- | --- | --- | --- | --- | --- | --- |
|  | 1 | 8 | 24 | 48 | 72 | 240 |
| **Hair** | 67% | 83% | 100% | 67% | 50% | 0% |
| Exudate | 83% | 83% | 17% | 0% | 0% | 0% |
| **Serum** | 100% | 83% | 67% | 50% | 33% | 0% |

### 3.4. Detection of *Loxosceles* DNA in rabbit samples by PCR

PCR detection could be observed for the hair and exudate samples being 8 h the time with the highest sensitivity (50 %) (Figure 4). It was not possible to confirm envenomation by PCR in any of the samples for the serum samples, which may indicate that the amount of DNA in the samples could be lower than the limit achieved with the technique (20 pg).

**Figure 4.**
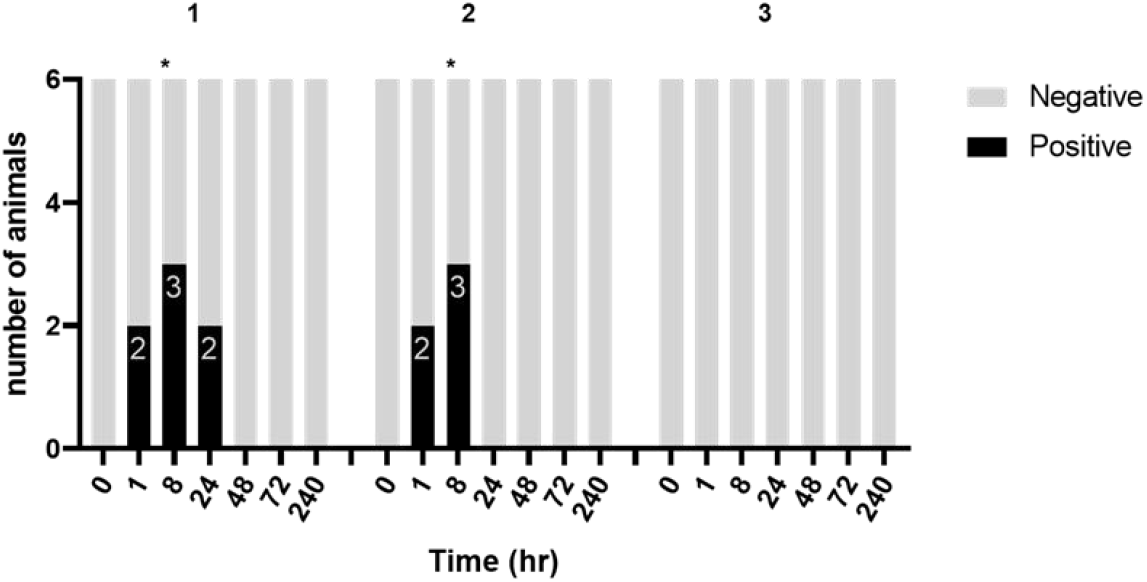
Summary of PCR test results for hair, exudate, and serum samples at different times after experimental rabbit envenomation. 1) Hair samples. 2) Exudate samples. 3) Serum samples. The asterisks above the columns indicate a statistically significant difference between animals before venom injection (time 0). The statistical parameters were calculated with the nonparametric chi square test.

## Discussion

Loop-mediated isothermal amplification (LAMP) is considered a tool with high specificity, sensitivity, simple, and quick execution, and with the possibility of visible results observation, thus meeting most of the requirements to categorize it as a point-of-care diagnostic [27].

We used in this work, as primers proportions and concentrations (0.2 μM of F3/B3, 0.4 μM of LF/LB, and 1.6 μM of FIP/BIP) the same proportions established by the creators of the technique [28] and others [11,29,30]. The stringent temperature of 71 °C was chosen due to the consistent results and lack of spurious amplification, respecting the operating range of the enzyme temperature (60-72 °C) [31]. We achieved better amplification using 60 min incubation even knowing that good results can be obtained in short intervals as short as 15 min [28]. There are already strategies aiming to reduce incubation time by using guanidine chloride or the use of additional primers for regions on the opposite strands and upstream to the inner primers (FIP and BIP) [32–34]not used in this work but that can be used to improve the reaction time.

Since LAMP conditions were standardized using the *L. similis* DNA, we also observed a high sensitivity in this test, in the range of 0.315 pg.

Using samples from *L. intermedia* experimentally-envenomed rabbits, we observed a higher sensitivity in the first 24 h after the envenomation, in which it was possible to detect *Loxosceles* DNA in the wound hair and serum from 1 to 24 h with the sensitivity ranging from 67 to 100 %. Previous works investigating *Loxosceles* envenomation using ELISA achieved a sensitivity of around 60 % [35,36].

In addition, it is possible to improve the sensitivity of the test, since some works have even reported strategies to improve the sensitivity and specificity of LAMP tests. Among these strategies, we can mention the addition of DMSO, TMSO, glycerol, and betaine, which are denaturing agents, help in the separation of DNA strands and facilitate the hybridization of the primers.

The specificity of LAMP was evaluated with DNA from organisms that, in humans, cause signs, mainly lesions, similar to loxoscelism leading to the misdiagnosis and impair adequate treatment. With our results, it was possible to confirm that the primers used were specific for samples containing *Loxosceles* DNA, as they did not amplify any of the other genetic materials tested.

Different from what was previously studied, we proposed to evaluate *Loxosceles* DNA in samples from experimentally envenomed animals in this work. That said, this was a preliminary study to assess the possibility of identifying DNA from *Loxosceles* spiders instead of venom protein components or antibodies already evaluated in other studies [35].

Comparing the evaluation time, in previous studies, the detection of protein components of *Loxosceles* venom could be done for up to 21 days [36]. The discrepancy in the detection times of this work (up to 72 h) compared to the others can be the result of DNA degradation by the action of deoxyribonuclease proteins released during the necrotic process resulting from the action of components present in the venom of *Loxosceles* spiders [37,38]. The aim of the study was accomplished since the LAMP test was able to identify the experimental envenomation within 1 h for all samples evaluated.

In conclusion, this is the first report demonstrating the use of LAMP to detect DNA from *Loxosceles* envenomation. Nevertheless, further studies are required to improve this technique and determine whether it has clinical applicability. If high diagnostic accuracy is confirmed in human cohorts, this method will be a valuable reference diagnostic tool for epidemiological investigations and clinical studies for brown spiders and other venomous animal envenomation.

## Author Contributions

L. P.F. performed experiments, generated all the data and figures. L.F.F: Conceived the research; C.G.D.: provided *L. similis* species; J.C.M: provided *Loxosceles* venoms; L.P.F, M. N.R., R.L.M.N and L.F.F discussed and analyzed the results, wrote the paper.

## Acknowledgments

Lucas Silva for graphical abstract design and SynBiom group for fruitful discussions. FUNED and FIOCRUZ for DNA control samples.

## Supplementary material

**Supplementary Figure S1:**
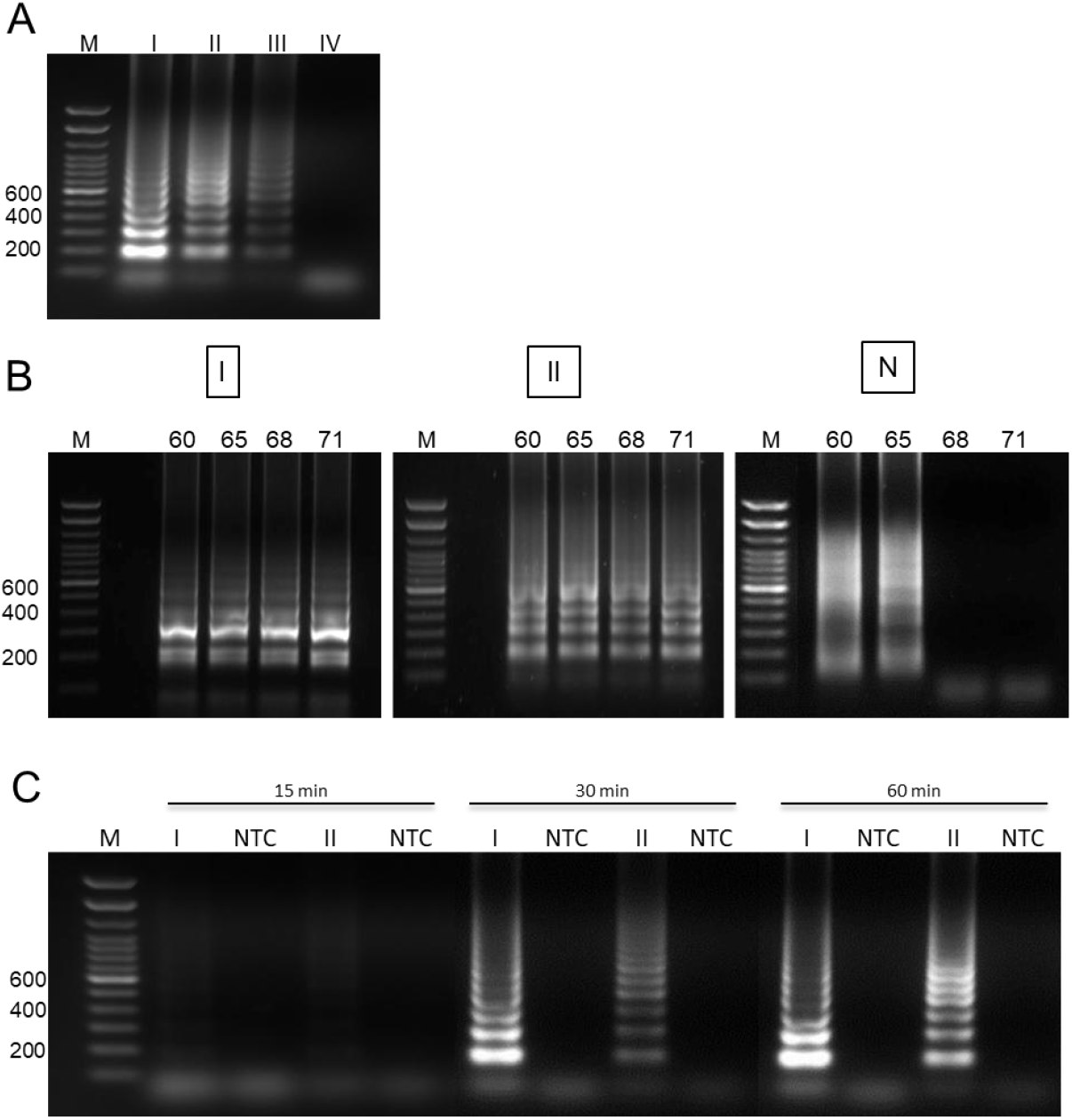
LAMP conditions evaluation. A) Different primer concentrations: I) F3/B3 0.2 uM, LR/LF 0.4 uM, FIP/BIP 1.6 uM; II)F3/B3 0.1 uM, LR/LF 0.2 uM, FIP/BIP 0.8 uM; III)F3/B3 0.05 uM, LR/LF 0.1 uM, FIP/BIP 0.4 uM; IV) NTC. All reactions were conducted using 10 ng of *L. similis* DNA. B) Different temperature conditions (60, 65, 68 and 71 °C) using primers conditions I, II. NTC: no template control. C) Different reaction time tested (15, 30 and 60 min) with 2 different primer conditions I and II. M: molecular weight.

**Supplementary Figure S2:**
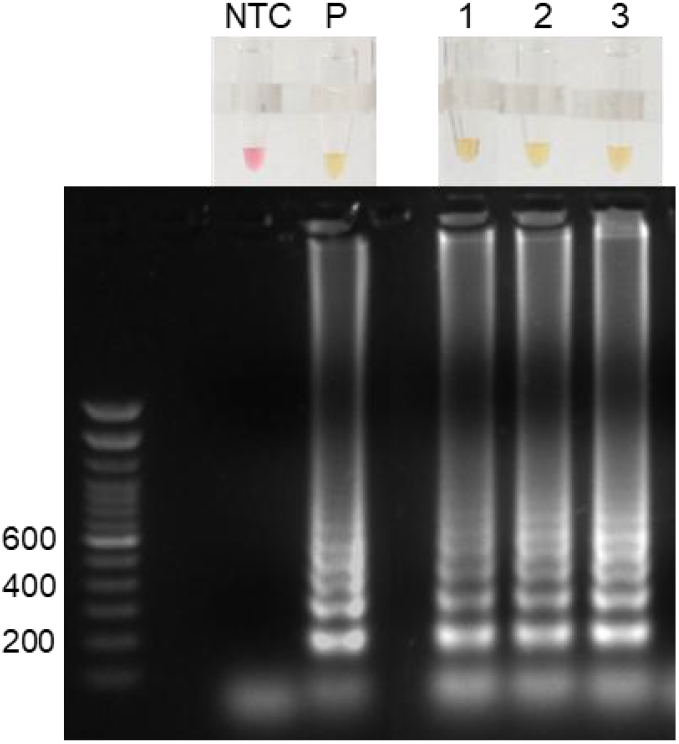
DNA detection of *Loxosceles intermedia* (*1*), *Loxosceles laeta* (*2*) and *Loxosceles gaucho* (*3*) in the raw venom. The DNA was extracted from 2 μL of the venom and used in the assay. NTC: Negative control; P: Positive control (2 ng of *L. similis DNA*).

